# Biodistribution and Environmental Safety of a Live-attenuated YF17D-vectored SARS-CoV-2 Vaccine Candidate

**DOI:** 10.1101/2022.01.24.477505

**Authors:** Li-Hsin Li, Laurens Liesenborghs, Lanjiao Wang, Marleen Lox, Michael Bright Yakass, Sander Jansen, Ana Lucia Rosales Rosas, Xin Zhang, Hendrik Jan Thibaut, Dirk Teuwen, Johan Neyts, Leen Delang, Kai Dallmeier

**Author notes:** equal contribution (Laurens Liesenborghs, Lanjiao Wang).

## Abstract

New platforms are urgently needed for the design of novel prophylactic vaccines and advanced immune therapies. Live-attenuated yellow fever vaccine YF17D serves as vector for several licensed vaccines and platform for novel vaccine candidates. Based on YF17D, we developed YF-S0 as exceptionally potent COVID-19 vaccine candidate. However, use of such live RNA virus vaccines raises safety concerns, *i*.*e*., adverse events linked to original YF17D (yellow fever vaccine-associated neurotropic; YEL-AND, and viscerotropic disease; YEL-AVD). In this study, we investigated the biodistribution and shedding of YF-S0 in hamsters. Likewise, we introduced hamsters deficient in STAT2 signaling as new preclinical model of YEL-AND/AVD. Compared to parental YF17D, YF-S0 showed an improved safety with limited dissemination to brain and visceral tissues, absent or low viremia, and no shedding of infectious virus. Considering yellow fever virus is transmitted by *Aedes* mosquitoes, any inadvertent exposure to the live recombinant vector via mosquito bites is to be excluded. The transmission risk of YF-S0 was hence evaluated in comparison to readily transmitting YFV-Asibi strain and non-transmitting YF17D vaccine, with no evidence for productive infection of vector mosquitoes. The overall favorable safety profile of YF-S0 is expected to translate to other novel vaccines that are based on the same YF17D platform.

## INTRODUCTION

Roughly two years after first emergence in 2019/2020, more than 5 million people have succumbed to Coronavirus Disease 2019 (COVID-19) caused by Severe Acute Respiratory Syndrome Coronavirus 2 (SARS CoV 2) (https://coronavirus.jhu.edu/map.html). Mass immunization is key to mitigating the expanding pandemic [1]. A set of rapidly developed prophylactic vaccines plays a crucial role in global immunization against SARS-CoV-2. Several of these vaccines are first-in-class based on novel platforms, including game changer mRNA vaccines and viral vector vaccines that are unprecedented in both, their high clinical efficacy as well as the incremental advance in breakthrough innovation [2-4]. However, a global vaccine supply shortage, the dependence on an ultra-cold chain system in case of mRNA vaccines, and the continuous emergence of virus variants pose unmet challenges [5, 6]. Unfortunately, long-term effectiveness of current SARS-CoV-2 vaccines is waning due to the combined effect of (*i*) a rapid decay of virus-neutralizing antibodies (nAb) over time and (*ii*) emergence of new variants escaping vaccine-induced immunity [7-9]. Furthermore, several first-generation COVID-19 vaccines have a rather high reactogenicity. With the growing number of vaccinated people, more cases and a wider spectrum of adverse effects following immunization (AEFI), including severe adverse effects (SAE) such a myocarditis or life-threatening deep-venous thrombosis are described [10-15]. In summary, there is an urgent to develop new and improved second-generation COVID-19 vaccines to quench the pandemic.

Recently, we used an alternative vaccine platform that uses the fully replication competent live-attenuated yellow fever vaccine YF17D as vector [16] and developed a virus-vectored SARS-CoV-2 vaccine candidate (YF-S0) that expresses a stabilized prefusion form of SARS-CoV-2 spike protein (S0) [17]. YF-S0 was shown to induce vigorous humoral and cellular immune responses in hamsters (*Mesocricetus auratus*), mice (*Mus musculus*) and cynomolgus macaques (*Macaca fascicularis*) and was able to prevent COVID19-like disease after single-dose vaccination in a stringent hamster model. Due to its YF17D backbone, YF-S0 could serve as dual vaccine to also prevent yellow fever virus (YFV) infections, which should provide an added benefit for populations living in regions at risk of YFV outbreaks [18].

In addition to preclinical efficacy, development of such a new vaccine requires in-depth evaluations of its safety to support progression from preclinical study to clinical trials. In particular for live-attenuated viral vaccines such as YF-S0, the biodistribution of the vaccine virus after administration needs to be assessed [19] to understand the viral organ tropism and hence to exclude potential direct harm to specific tissues. Our vaccine candidate YF-S0 showed an excellent safety profile in multiple preclinical models, including in NHP as well as in interferon-deficient mice and hamsters [17]. However, use of such a recombinant YF17D vaccine entails some potential concerns [19]. Particularly, replication and persistence of YF-S0 in tissues and body fluids poses a theoretical risk of YF vaccine-associated viscerotropic disease (YEL-AVD) and YF vaccine-associated neurotropic disease (YEL-AND), which are originally linked to parental YF17D [20]. Regarding to this, the parental YF17D vaccine are commonly used as benchmark for direct comparison in safety assessment [19].

Here, we investigated the biodistribution and shedding of YF-S0 following vaccination in hamsters, with as aim to understand (*i*) to what extent YF-S0 causes viremia resulting into virus dissemination to vital organs; (*ii*) to evaluate the risks of YF-S0 for YEL-AVD/AND by confirming its transient and self-limited replication *in vivo* [17], restricting the risks for YEL-AVD/AND; (*iii*) to what extent viral RNA remains detectable in body secretions and, in case, (*iv*) if this poses any environment risks for shedding of recombinant infectious virus. Furthermore, YFV is also a mosquito-borne virus. To eliminate the concerns that YF-S0, which employs licensed YF17D as a vector and hence, despite being proven highly attenuated, might lead to an increased environmental risk causing by phenotypical change as any other recombinant viruses. Taking this theoretical consideration into account, we tested the infectivity of YF-S0 on *Aedes aegypti* (*Ae. aegypti*) mosquitoes to assess its transmission potential. *Ae. aegypti* was selected as target mosquito species because of its well-known high vector competence for YFV [21]. It is well documented that wild-type YF-Asibi can infect and disseminate in *Ae. aegypti* while YF17D only occasionally infects the midgut and is unable to disseminate to secondary organs [22, 23]. Therefore, these two YFV strains were used as controls to assess transmission of YF-S0 by a competent vector.

Finally, we corroborate the favorable safety profile of YF-S0 by reporting limited dissemination and shedding in vaccinated hamsters, nor any risk of mosquito-borne transmission.

## RESULTS

### Tissue distribution of YF-S0 and parental YF17D in hamsters

For our assessment, we chose wild-type (WT) Syrian golden hamsters as preferred small animal model of YFV infection [24] and injected them with a high dose (10^4^ PFU) of either YF17D (n=6) or YF-S0 (n=6) via intraperitoneal route to achieve maximal exposure; with primary pharmacodynamics documented before [17] and confirmed here by consistently high seroconversion rates (at least 80%) to YFV-specific nAb (Fig. S1). As methods control, we inoculated STAT2-knockout (STAT2^-/-^) hamsters with 10^4^ PFU of YF17D (n=2). STAT2^-/-^ hamsters are deficient in antiviral type I and type III interferon responses [25] and therefore prone to uncontrolled flavivirus replication [26]. Tissues sampled for analysis were chosen based on biodistribution data available from non-human primates and humans. In macaques, detection of YF17D RNA has been reported in lymph nodes, spleen and liver at 7 days post subcutaneous inoculation [27]. Likewise, viral RNA is widespread and abundantly found in spleen, liver, brain, kidney, and other organs in patients who developed YEL-AVD [20, 28]. Based on this knowledge, we collected spleen, liver, brain, and kidney as most common target organs to assess the risks for YEL-AVD and YEL-AND. Ileum and parotid gland were collected as additional excretory tissues, and lung as main target of COVID19 (Fig. 1A). From our previous experience [17], we observed that the replication of YF17D or YF-S0 is transient and well tolerated in WT hamsters. Tissue analysis in hamsters was thus performed 7 days post inoculation (dpi), *i*.*e*., few days after peak of viremia and at a timepoint which STAT2^-/-^ hamsters needed to be euthanized for humane reasons.

**Fig. 1.**
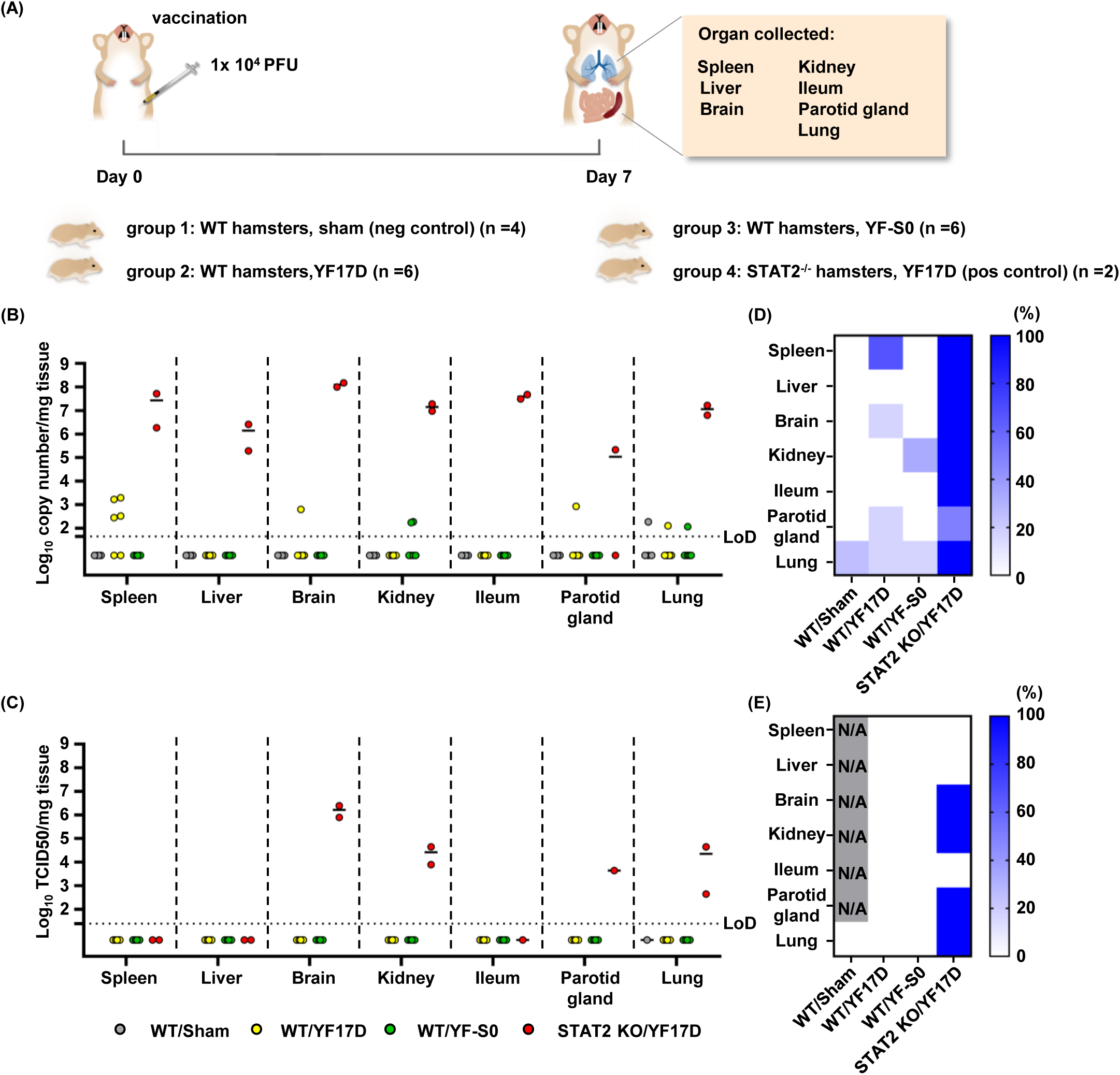
Biodistribution of YF-S0 in hamsters. **(A)** Schematic of hamster vaccination and organ collection. Hamsters were inoculated i.p. with 104 pfu/ml of either YF17D or YF-S0 and sacrificed 7 days later. Organs from 4 different experimental groups, including Sham vaccinated wild-type (WT) hamsters and YF17D vaccinated STAT2^-/-^ knock out (KO) hamsters as respective negative and positive controls, were collected and divided for RNA extraction and virus isolation. **(B)** Viral RNA load by RT-qPCR. (**C**) Virus isolation by TCID50 assay. For Sham and STAT2 KO, only PCR-positive samples were analyzed. **(D**,**E)** Heat map representing positivity rates by organ and experimental group based on the results of RT-qPCR (D) or TCID50 assay **(E)**. Bars in **(B)** and **(C)** represent median values. N/A: not applicable

Viral RNA above detection limits in YF17D vaccinated WT hamsters was mostly limited to spleen (4/6), with exception of a single hamster in which viral RNA was widespread to brain, parotid gland, and lung (Fig. 1B and Suppl Table 1). Detection of YF-S0 was markedly less frequent and restricted to only kidney (2/6) and lung (1/6) (Fig. 1B and Fig. 1D). Overall, in either group RNA level was low and barely detectable by sensitive RT-qPCR, indicative for limited replication in WT hamsters. In contrast, unrestricted replication of virus to high viral loads was observed in STAT2^−/−^ hamsters (Fig. 1B and Fig. 1C). Importantly, no viral RNA nor infectious virus could be detected in brains of YF-S0 vaccinated hamsters, suggesting a low associated YEL-AND risk (Fig. 1D and Fig. 1E).

Viremia is considered a key indicator for the risk of developing YEL-AVD. Kinetics of viral RNA in serum as proxy for viremia have been reported earlier for WT hamsters vaccinated with YF17D or YF-S0 [17] and are discussed here in comparison to respective data from STAT2^-/-^ controls (Fig. 2B). Viremia can be detected consistently in all YF17D vaccinated WT hamsters (6/6) starting at 1 dpi and lasting for 2.5 (1-4) days in median (95% confidence interval); by contrast, viral RNA was detected only once at 3 dpi in a single YF-S0 vaccinated hamster (1/6) (Fig. 2B and Suppl. Table 2). In STAT2^-/-^ hamsters, YF17D grew unrestrictedly to markedly increased viral RNA levels (Fig. 2B), readily detectable by virus isolation (Fig. S2). Integration of data over the course of immunization (area under the curve, AUC) indicated a significant reduced overall serum virus load in YF-S0 vaccinated animals (Fig. 2F).

**Fig. 2.**
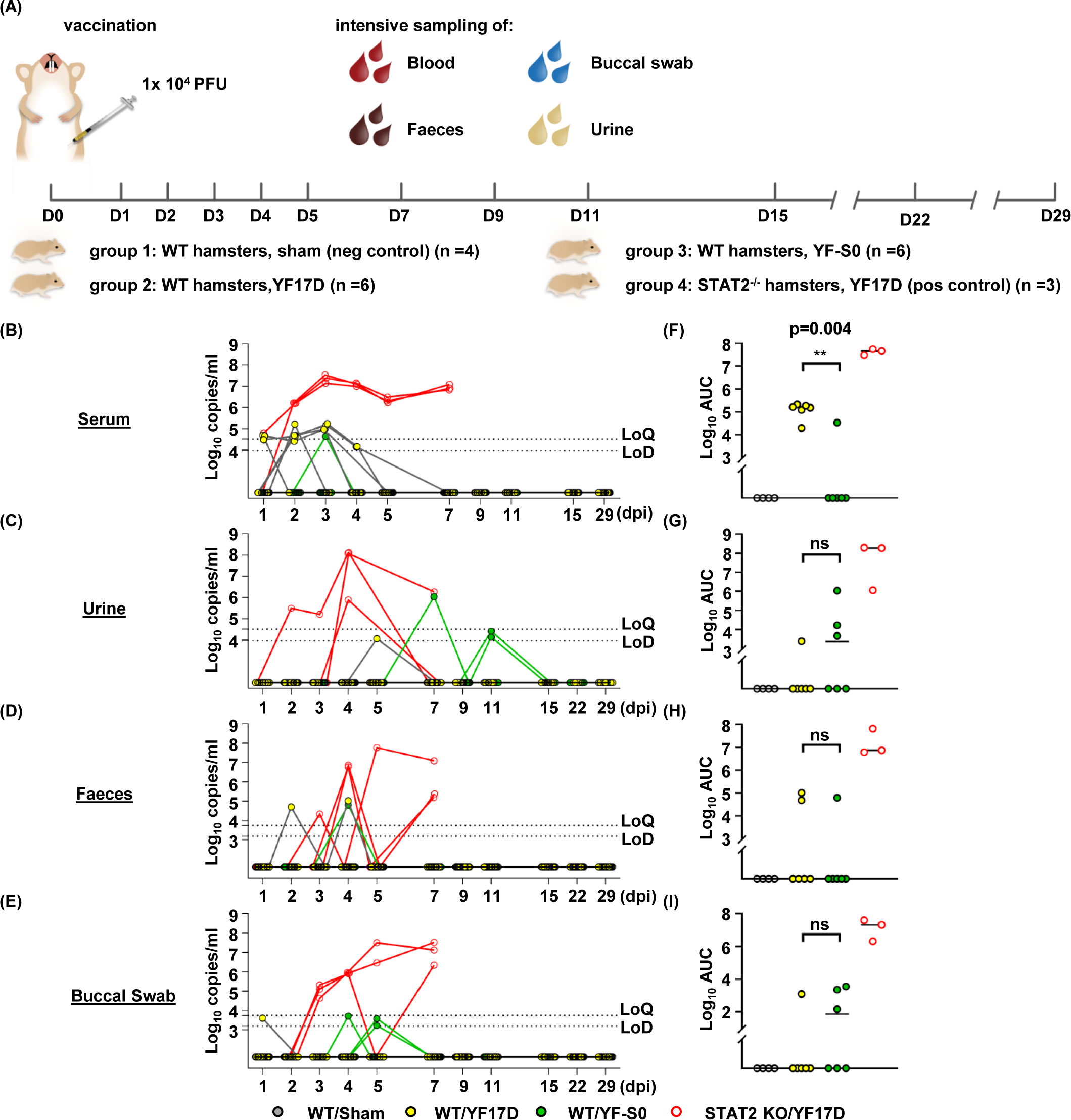
Shedding of YF-S0 by vaccinated hamsters. **(A)** Schematic of vaccination and specimens’ collection. Hamsters were inoculated as in Fig. 1A, and serum, urine, faeces, and buccal swabs serially sampled at indicated timepoints. **(B)-(E)** Viral RNA load by RT-qPCR. **(F)-(I)** Area under curve (AUC, copies*day) calculated by GraphPad Prism 8; Mann-Whitney test was used for the statistic analysis, with p >0.05 marked as non-significant (ns), and p ≤ 0.01 as **. Serum RNA data for YF17D and YF-S0 vaccinated WT hamsters as previously published. LoQ: Limit of quantification; LoD: Limit of detection; dpi: days post inoculation.

### Limited shedding of YF-S0 and parental YF17D RNA

Since shedding of viral RNA in urine after YF17D vaccination has been reported [29], we sampled different body fluids to investigate respective virus levels (Fig. 2A). Within all longitudinally sampled specimens, viral RNA was detected only sporadically in urine (1/56; 3/58), faeces (2/65; 1/66), and buccal swabs (1/66; 3/66) of both YF17D and YF-S0 vaccinated hamsters, mostly at very low copy numbers, and not linked to viremia (Figure 2C-2E, Fig. S3, and Suppl Table 3-5). Noteworthy, viral RNA could only be detected, if at all, only within the first 11 dpi, clearly indicating that viral replication was self-limiting, leading to the final elimination of the live viral vector from all tissues. Also, there was no significant difference regarding the AUC between both groups (Fig. 2G-2I). The potential risk of YF-S0 to be spread by excrements of vaccinated individuals should hence be as low as for YF17D. In addition, no viable virus could be isolated from urine samples with RNA counts as high as 10^8^ copies/mL (not shown), in line with no clinical evidence for secondary spread in urine, matching longstanding field experience for YF17D.

### Abortive infection of YF-S0 on yellow fever virus competent vector *Ae. aegypti*

YF-S0 is derived from mosquito-borne YFV, and human-to-human transmission by a competent mosquito vector could theoretically lead to unintentional exposure to the vaccine, including in immune-compromised people [30]. Thus, the transmission risk of YF-S0 should be excluded regarding main indicators of mosquito vector competence [21, 31, 32] (Fig. 3A): (*i*) sufficient virus ingestion from infectious blood meal; (*ii*) productive infection of virus in mosquito midgut (midgut infection barrier, MIB); and, (*iii*) virus escapes from midgut barrier (MEB), *i*.*e*., dissemination to parenteral tissues to establish sufficiently high virus loads in salivary glands to enable transmission. To this end, *Aedes aegypti* mosquitoes, as the species of YFV competent vector [21], were given infectious blood meals with either no virus, YF17D, YF-S0 or wild-type YF-Asibi strain as positive control [22, 23]. Infection was determined by RT-qPCR and virus isolation on day 0 on whole mosquitoes (ingestion step), on day 14 in thorax and abdomen (virus infection and replication in mosquito midgut; marked as main body), and on day 14 dissemination in head, legs and wings (dissemination).

**Fig. 3.**
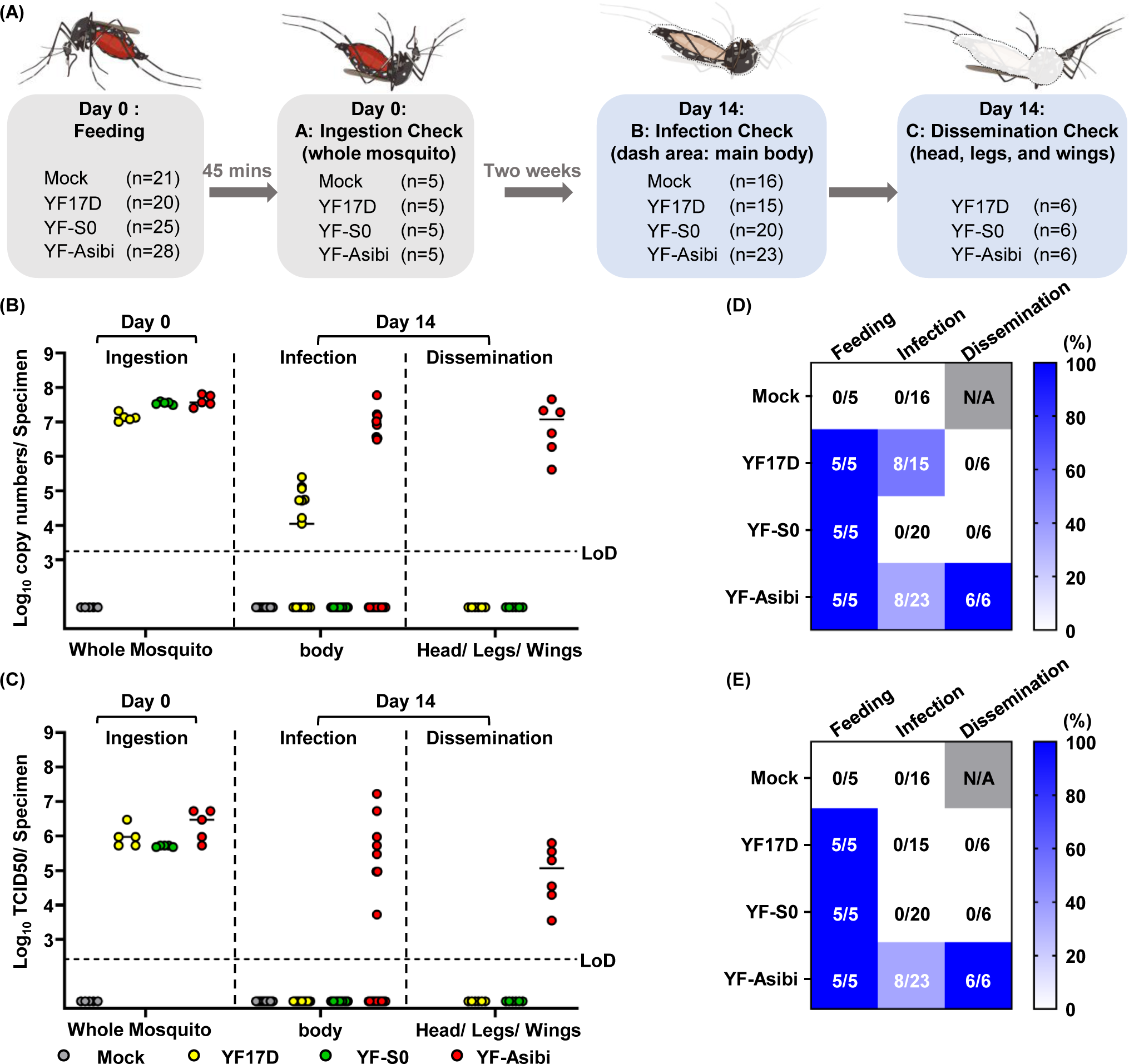
Assessment of YF-S0 transmission potential by Aedes mosquitoes. **(A)** Schematic of virus feeding of mosquitoes and specimens’ collection. Mosquitoes were fed with infectious blood meal containing YF17D, YF-S0 or YF-Asibi, or Mock. 5 mosquitoes were collected each for ingestion assessment. At 14 days post feeding (dpf), remaining mosquitoes were dissected into two parts, midgut (infection assessment) and head, legs, and wings (dissemination assessment). **(B)** Viral RNA load by RT-qPCR. **(C)** Virus isolation by TCID50 assay. For assessment of ingestion and infection, RT-qPCR and TCID50 were performed on all samples. For assessment of dissemination, only a selection of PCR-positive specimens from the YF17D and YF-Asibi groups (n=6 each) were further analyzed by TCID50 assay, plus 6 randomly chosen from the YF-S0 group. **(D&E)** Heat map representing positivity rates per experiment group as scored by RT-qPCR **(D)** and TCID50 assay **(E)**. Bars in **(B)** and **(C)** represent median values. N/A: not applicable. Mosquito icons were adapted from BioRender.com (2021).

Experimental feeding was equally efficient for all three virus groups regarding both viral RNA and infectious virus recovered (Fig. 3B&C). However, 14 days after feeding, viral RNA was detected exclusively in specimens from the YF17D group (8/15) and YF-Asibi group (8/23); yet none from the YF-S0 group. Importantly, infectious viral particles were only detectable in the YF-Asibi group, with virus loads as high as about 10^6^ TCID_50_/body on average (Fig. 3C). For dissemination beyond the MEB, the remaining head, legs and wings of each six virus-positive mosquitoes with highest body virus loads from the YF17D and YF-Asibi groups, respectively, and six randomly chosen specimens from the YF-S0 group were evaluated. All these specimens from the YF-Asibi group (6/6) scored positive for dissemination, while none from the YF-S0 or YF17D groups (Fig. 3B-C). The results showing that YF-S0 is neither able to pass the MIB for midgut infection, nor to escape from the midgut (MEB) for dissemination (Fig. 3D&E).

## DISCUSSION

The live-attenuated YF17D vaccine is considered as one of the most powerful and successful vaccines and has been used on humans for decades [33]. Its well-known characteristics of stimulating both vigorous humoral and cellular immune responses, as well as favorable innate responses is of interest for other vaccine targets using the YF17D genome as a backbone [16]. We recently generated a particularly potent YF17D-vectored vaccine candidate, YF-S0, against SARS-CoV-2 infection, inserting the non-cleavable spike protein of SARS-CoV-2 (S0) between the E and NS1 region of YF17D [17]. This construct serves as antigens to induce vigorous immune responses against both SARS-CoV-2 and YFV infections [17].

Apart from YF-S0, YF17D is currently the only fully replication competent viral vector that is part of any licensed recombinant live viral vaccine in wide use for human medicine; *i*.*e*. in the two licensed human vaccines, JE-CV (against Japanese encepahilitis; Imojev® [34]) and CYD-TDV (against all four serotypes of dengue virus; Dengvaxia® [35]). Additional YF17D-based vaccine candidates are in different stages of (pre)clinical developed, including vaccines against other flaviviruses (West Nile virus: ChimeriVax-WN02 [36]; Zika virus: YF-ZIKprM/E [37]) or non-flaviviruses (HIV: rYF17D/SIVGag45–269 [38]; Lassa virus: YFV17D/LASVGPC [39]; chronic hepatitis B virus: YF17D/HBc-C [40]). As these YF17D-vectored vaccines has been proved, YF-S0 also trigger vigorous protective immune responses, including high levels of SARS-CoV-2 neutralizing antibodies after a single dose vaccination in hamsters, mice and cynomolgus macaques. However, despite little (pre)clinical evidence nor such reports from post-marketing surveillance, all these YF17D-vectored vaccines share the theoretical concerns of the SAEs associated with the parental YF17D vaccines, such as YEL-AVD (0.4 per 100,000) and YEL-AND (0.8 per 100,000) [19, 20, 41, 42].

To temper the remaining safety concerns, the viscerotropism and neurovirulence of YF-S0 was compared head-to-head with parental YF17D virus by investigating the biodistribution and viremia following administration of either vaccine virus in hamsters. We demonstrate that parental YF17D can spread systemically and viral RNA can be detected in spleen, brain, parotid gland, and lung in YF17D vaccinated WT hamsters. However, replication of YF17D remains restricted, resulting in infectious virus loads below detection limits. Compared to YF17D, detection of YF-S0 was further limited, with minute amounts of viral RNA in kidney and lung. Unrestricted virus replication to high viral loads as cause of viscerotropic or neurotropic disease was observed only in STAT2^−/−^ hamsters, in line with the essential role innate interferon signaling plays in live vaccines [30, 43] and control of viral infections in general [44]. In addition, in YF-S0 vaccinated WT hamsters, detection of viremia was rare (Fig. 2B) and importantly, less frequent (1/6) and markedly lower in magnitude (AUC) and duration (1 day) compared to parental YF17D (6/6 for >2 days). Taken together, the overall limited tissue distribution of YF-S0 as well as the low abundance of its RNA in blood, below detection limits for infectious virus, suggest a further lowered risk of YEL-AVD/AND for YF-S0 than that reported parental YF17D. To further investigate the potential environment risk associated with shedding of recombinant virus, we collected urine, faeces and buccal swabs from vaccinated hamsters and checked for the presence of viral RNA for 29 days to determine how long YF-S0 would remain detectable in body secretions as compared to YF17D. No significant differences in vaccine RNA shedding were observed between YF17D and YF-S0 during the course of immunization (Fig. 2G-2I, AUC). Importantly, no infectious virus could be isolated, suggesting the risk is very low, even if any inadvertent exposure by vaccinated individuals to their environment. In summary, these results obtained in a hamster model of YF17D vaccination clearly demonstrate that (*i*) the overall viral tissue burden for YF-S0 was considerably lower than for parental YF17D, and (*ii*) presence of viral RNA in body secretions (urine, feces, and buccal swab) was equally low as for YF17D, mostly likely void of residual infectious virus particles. YF-S0 vaccine virus infection is transient and harbors minimal, if at all any, risk of shedding nor evidence for environmental biosafety concern.

Last, though the chances of YF17D-vectored vaccines to be transmitted by arthropod vectors are minimal, we evaluated the replication competence of YF-S0 in yellow fever mosquito vector (i.e., *Ae. aegypti*). While parental YF17D passed the MIB and got restricted at the MEB as previous documented [22, 23], YF-S0 was already blocked at the first barrier with no remaining viral RNA or infectious virus detectable after an infectious bloodmeal. Hence, the transmissibility of YF-S0 by mosquitoes is to be considered neglectable.

Altogether, YF-S0 is considered a safe and efficacious vaccine candidate for the prevention of COVID19. A similar improved safety as compared to parental YF17D can be expected for other vaccines following the same design principle, *i*.*e*., using transgenic, yet fully replication-competent YF17D as vector [16, 40].

## Funding

This project has received funding from the European Union’s Horizon 2020 Research and Innovation Program under grant agreement no. 733176 (RABYD-VAX to K.D. and J.N.), and no. 101003627 (SCORE to J.N.). Funding was further provided by the Research Foundation Flanders (FWO) under the Excellence of Science (EOS) program (no. 30981113, VirEOS to K.D and J.N.), the FWO COVID19 call (no. G0G4820N), and by the KU Leuven/UZ Leuven COVID-19 Fund (COVAX-PREC project). L.H.L. acknowledges support by a PhD scholarship grant from the KU Leuven Special Research Fund (DBOF/14/062). L.L. is member of the Institute of Tropical Medicine’s Outbreak Research Team which is financially supported by the Department of Economy, Science and Innovation (EWI) of the Flemish Government. X.Z. was supported by a PhD scholarship grant from the China Scholarship Council (CSC, no. 201906170033). L.D. received funding from KU Leuven Internal Funds (C22/18/007 and STG/19/008), as well as K.D. (C3/19/057 Lab of Excellence). This publication was supported by the Infravec2 project, which has received funding from the EU’s Horizon 2020 Research and Innovation Program 2020 (grant agreement no. 731060).

## Contributions

Overall conceptual design: L.-H.L., L.L., D.T., and K.D.; methodology: L.-H.L., L.L., L.W.; in vivo experiments (hamsters): L.L. and M.L.; in vivo experiments (mosquitoes): L.W. and A.L.R.R.; in vitro experiments: L.-H.L. and X.Z.; in vitro experiment (serology): H.J.T.; design and generating of YF-Asibi: M.B.Y. and S.J.; data management and analysis: L.-H.L. and L.L.; writing of manuscript-draft: L.-H.L.; writing of manuscript-review and editing: L.-H.L., L.L., L.W., L.D., D.T., and K.D.; supervision: J.N. and K.D.; funding acquisition: J.N. and K.D.

## Acknowledgements

We thank Thibault Francken, Dagmar Buyst, Niels Cremers, Birgit Voeten, and Jasper Rymenants for their technical support on specimens’ preparation and virus titration as well as Jasmine Paulissen for her technical assistance on generating serology data.

## Data availability

All data supporting the findings in this study are available from the corresponding author upon request.

## Competing interests

The authors declare that there are no competing interests. This manuscript is currently under peer review.

## Materials and Methods

### Animal experiment

#### Hamsters

Wild-type (WT) outbred specific pathogen-free Syrian hamsters (*Mesocricetus auratus*) were purchased from Janvier Laboratories, France. The generation [45] and characterization [25] of STAT2^−/−^ (gene identifier: 101830537) hamsters has been described elsewhere. STAT2^−/−^ hamsters were bred in-house. Hamsters (max. n=2) were housed in individually ventilated cages (Sealsafe Plus, Tecniplast; cage type GR900), under standard conditions of 21 °C, 55% humidity and 12:12 light:dark cycles. Hamsters were provided with food and water ad libitum, as well as extra bedding material and wooden gnawing blocks for enrichment as previously described. This project was approved by the KU Leuven ethical committee (P015-2020), following institutional guidelines approved by the Federation of European Laboratory Animal Science Associations (FELASA). Hamsters were euthanized by intraperitoneal administration of 500 μL (hamsters) Dolethal (200 mg/mL sodium pentobarbital, Vétoquinol SA).

#### Vaccine and virus stocks

Vaccine viruses used throughout this study have been described [17]. YF-S0 was derived from a cDNA clone of YF17D (GenBank: X03700) with an in-frame insertion of a non-cleavable version of the SARS-CoV-2 S protein (GenBank: MN908947.3) in the YFV E/NS1 intergenic region. YF-S0 vaccine stocks were grown on baby hamster kidney (BHK21) cells. The molecular and antigenic structure and replication of YF-S0 has been described in detail [17]. Original YF17D vaccine (Stamaril, Sanofi-Pasteur; lot number G5400) was purchased via the pharmacy of the University Hospital Leuven and passaged twice in Vero E6 cells prior to use. The construction and rescue of YF-Asibi from an infectious cDNA clone will be described elsewhere (Yakass, Jansen et al.). The respective YFV cDNA sequence was adjusted to match previously described molecular clone Ap7M [46]. All virus stocks were titrated by plaque assay on BHK21 cells [17].

#### Biodistribution

WT hamsters (6-8 weeks old, female) were inoculated intraperitoneally with 10^4^ PFU/mL dose of YF17D (n = 6) or YF-S0 (n = 6). STAT2^−/−^ hamster (6-8 weeks old, female) were inoculated intraperitoneally with 10^4^ PFU/mL of YF17D (n = 2). At 7 dpi, blood, spleen, liver, brain, kidney, ileum, parotid gland, and lung were collected.

#### Shedding

WT hamsters (6-8 weeks old, female) were inoculated intraperitoneally with 10^4^ PFU/mL of YF17D (n = 6) or YF-S0 (n = 6). STAT2^−/−^ hamsters (6-8 weeks old, male) were inoculated intraperitoneally with 10^4^ PFU/mL of YF17D (n = 3). Blood, urine, faces, and buccal swab were collected daily for the first 5 dpi, then every other day until 11 dpi and 15, 22 (except for the blood) and 29 dpi, and afterwards once a week until 29 dpi.

### Mosquito experiment

#### Mosquito strain

*Ae. aegypti* Paea [47] were obtained via the Infravec2 consortium (*https://infravec2.eu/product/live-eggs-or-adult-females-of-aedes-aegypti-strain-paea-2/*) from Institute Pasteur of Paris. Mosquitoes were maintained at the insectary of Rega Institute, and the fourth generation was used for this study. In brief, larvae were fed with yeast tablets (Gayelord Hauser, France) until the pupae stage prior to transfer to cages for emergence. Adults were maintained with cotton soaked in 10% sucrose solution under standard conditions (28°C, 80% relative humidity, and 14h:10h light/dark cycle).

#### Oral infection and sample collection

7-day-old female mosquitoes were starved 24 h prior to infection. Infectious blood meals contained rabbit erythrocytes plus 5 mM adenosine triphosphate as phagostimulant, supplemented with virus stocks to final titers of 2×10^5^ PFU/mL for both YF17D and YF-S0, and 5×10^6^ PFU/mL for YF-Asibi, respectively. After 45 minutes, 5 full engorged females from each group were frozen for viral input assessment (ingestion check, Fig. 3A), and the rest kept with 10% of sugar solution under both controlled conditions (28 ± 1°C, relative humidity of 80%, light/dark cycle of 14h/10h, supplied with 10% sucrose solution) and BSL-3 containment conditions. At 14 dpi, mosquitoes were dissected into two parts; main body (thorax and abdomen) and remainder, collected individually in tubes containing PBS and 2.8 mm ceramic beads (Precellys). The samples were homogenized and pass through 0.8μm column filters (Sartorius, Germany). Thus, cleared supernatants were used for TCID_50_ assay or keep at −80°C for RNA extraction and subsequent RT-qPCR analysis.

#### RNA extractions

Solid tissues (organs), faeces and buccal swabs were homogenized in a bead mill (Precellys) in lysis buffer (Macherey-Nagel; cat no. 740984.10). After homogenization, samples were centrifuged at 10,000 rpm for 5 min to remove cell debris, and total RNA was extracted by using NucleoSpin Plus RNA virus Kit (Macherey-Nagel, cat no. 740984.10). For serum (50 μl), urine (50 μl) and homogenates of mosquito samples (150 μl), NucleoSpin RNA virus kit (Macherey-Nagel; cat no. 740956.250) was used for RNA extraction.

#### RT-qPCR

RT-qPCR for YFV detection was performed as previously described [17] using primers and probe targeting the YFV NS3 gene [23] on an ABI 7500 Fast Real-Time PCR System (Applied Biosystems). Absolute quantification was based on standard curves generated from 5-fold serial dilutions of YF17D cDNA with a known concentration.

#### TCID_50_ assay

For virus isolation and quantification BHK21 cells were infected with 10-fold serial dilutions in 96-well plates, and incubated at 37°C for 6 days using DMEM with 2% fetal bovine serum (Hyclone), 2 mM L-glutamine (Gibco), 1% sodium bicarbonate (Gibco), and 1% antibiotics (PenStrep) as assay medium. Solid tissues were homogenized in a bead mill (Precellys) in assay medium, and centrifuged at 10,000 rpm for 5 min (4°C) to remove debris. Resulting viral titers were calculated by the Reed and Muench method.

#### Serum neutralization test (SNT)

Titers of YFV-specific neutralizing antibodies were determined using BHK21 cells and a mCherry-tagged variant of YF17D virus (YFV-mCherry) as described [17]. In brief, YFV-mCherry was mixed and incubated with serial diluted of sera for 1 h at 37°C, and subsequently transferred to BHK21 cells grown in 96-well plates for infection. At 3 days post infection, the relative infection rate was quantified by counting mCherry-expressing cells versus total cells on a high content screening platform (CX5, Thermo Fischer Scientific), normalizing the infection rate of untreated virus controls as 100%. Half-maximal serum neutralizing titers (SNT_50_) were determined by curve fitting in GraphPad Prism 8.

#### Statistics

Data were analyzed using GraphPad Prism 8. Results are represented as individual values and median for summary statistics. Statistical significance was determined using non-parametric Mann–Whitney U-test (*P≤ 0.05; **P ≤ 0.01; ns, not significant)

---

**Fig. S1 (related to Fig. 1A and Fig. 2A).**
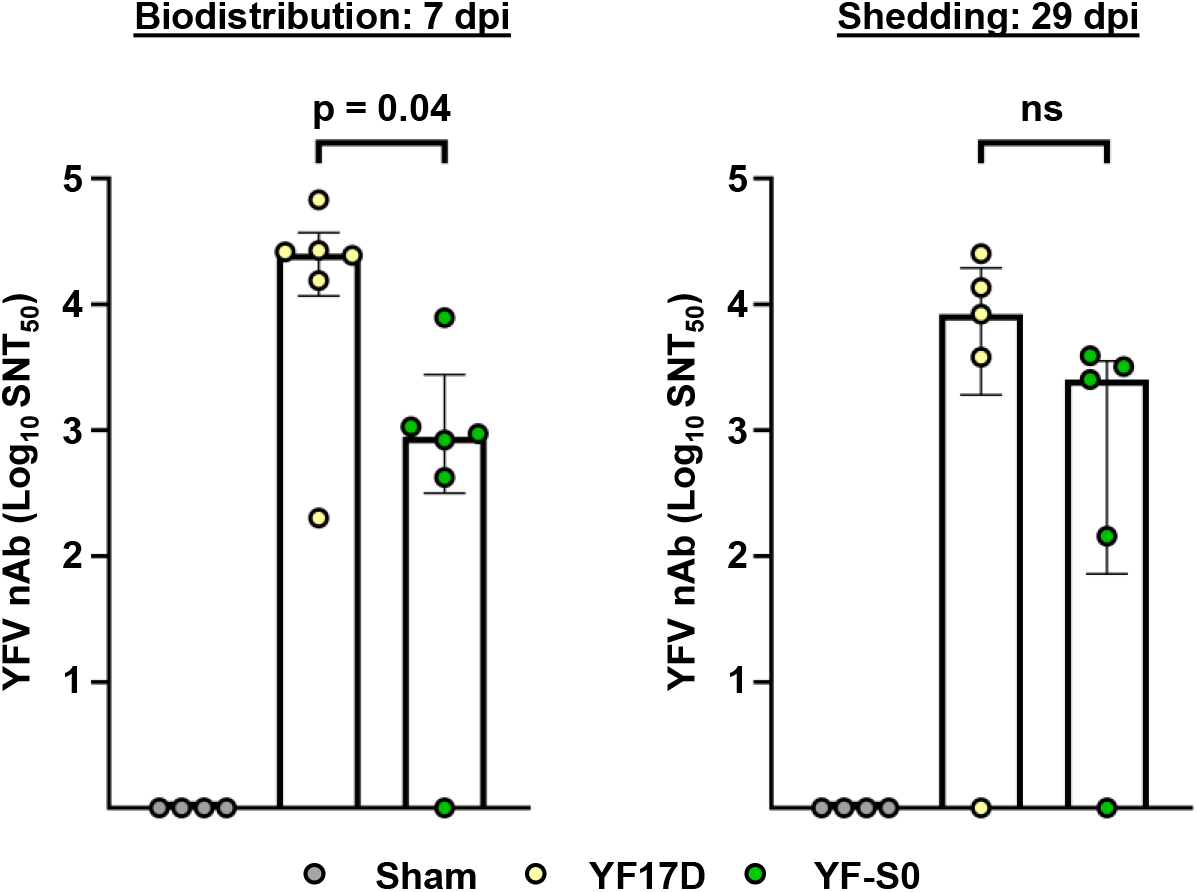
YFV-specific humoral immune responses in one-dose YF17D and YF-S0 vaccinated WT hamsters (primary pharmacodynamics). Serum samples for determination of YFV-specific neutralizing antibodies (nAb) collected at the respective endpoint of experiments assessing vaccine virus biodistribution (Fig.1A, 7 dpi) and shedding (Fig. 2A, 29 dpi). Sample number in biodistribution experiment: Sham n=4, YF17D n=6, and YF-S0 n=6, and in shedding experiment: Sham n=4, YF17D n=5, and YF-S0 n=5. 50% serum neutralizing titers (SNT_50_) were presented as median ± IQR for each group at logarithmic scale. Mann-Whitney test was used for the statistical analysis, with p >0.05 marked as non-significant (ns).

**Fig. S2 (related to Fig. 2A).**
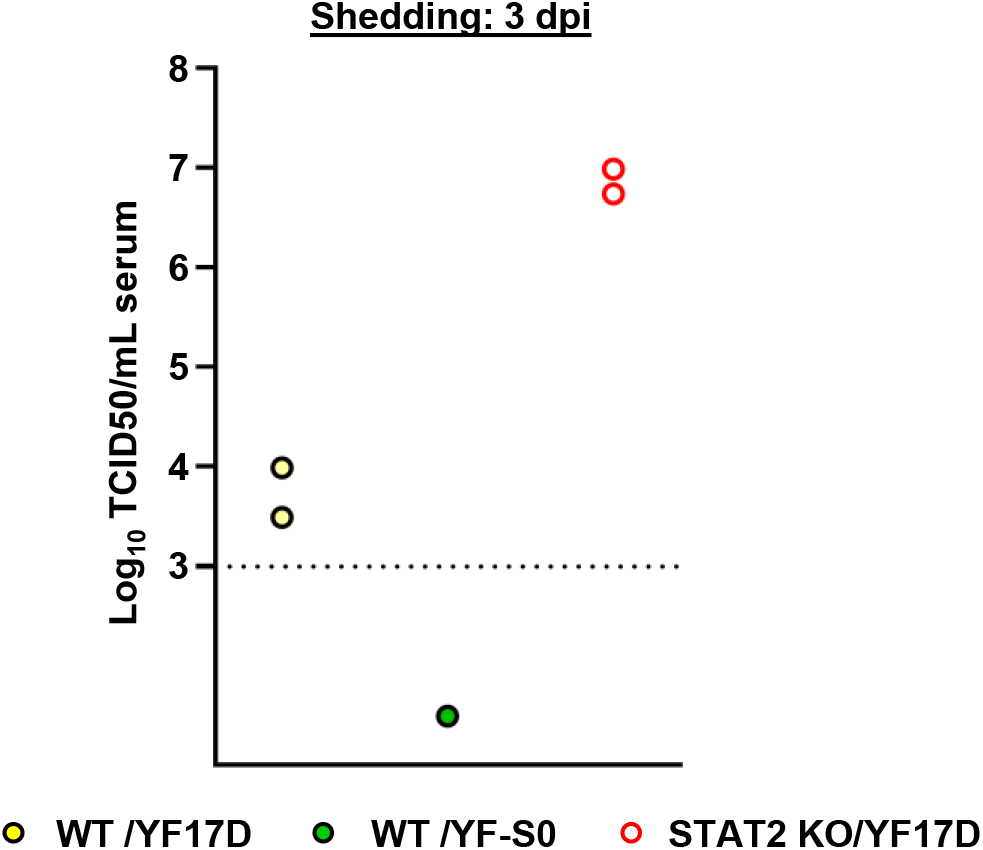
Infectious virus loads in serum (viremia) in selected YF17D or YF-S0 vaccinated hamsters. Serum samples collected at 3 days after vaccination. Sample number for YF17D vaccinated WT hamster, n=2; for YF-S0 vaccinated WT hamster, n=1; and (3) YF17D vaccinated STAT2^-/-^ hamster, n=2. Selected samples included specimen with respectively highest viral RNA copies numbers detected (YF17D in WT and STAT2^-/-^ hamsters) or, in case of YF-S0, the only PCR positive specimen.

**Fig. S3 (related to Fig. 2A).**
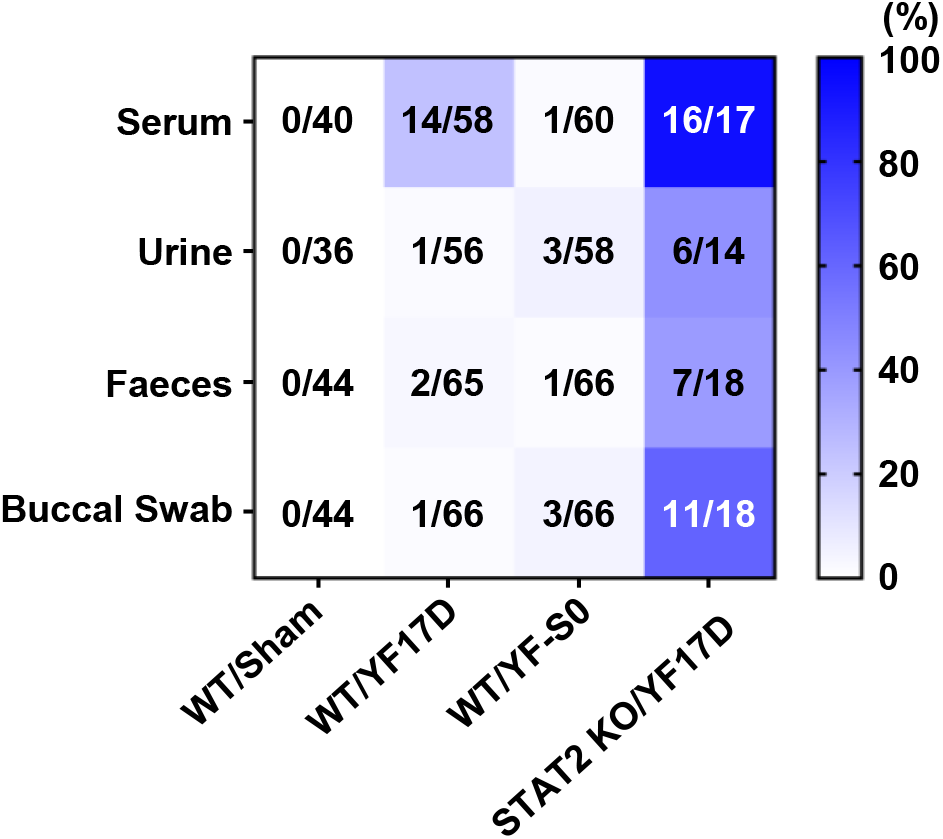
Cumulative detection rates of viral RNA by RT-qPCR in all specimens (serum, urine, faeces or buccal swab) collected from vaccinated WT or STAT2^-/-^ hamster. Specimens were collected according to sampling scheme depicted in Fig. 2A. Heatmap showing ratios of the total number of all PCR-positive samples versus the total number of all samples tested per study group over the course of 29 days after vaccination.

