## Supplementary Tables for "Biodistribution and Environmental Safety of a Live-attenuated YF17D-vectored SARS-CoV-2 Vaccine Candidate"

**Supplementary Table 1. (Related to Fig. 1B, RT-qPCR)**

| Sample | Genotype | vaccine | spleen | liver | brain | kidney | ileum | parotid gland | lung |
|---|---|---|---|---|---|---|---|---|---|
| 581 | WT | sham | n.d. | n.d. | n.d. | n.d. | n.d. | n.d. | n.d. |
| 582 | WT | sham | n.d. | n.d. | n.d. | n.d. | n.d. | n.d. | n.d. |
| 583 | WT | sham | n.d. | n.d. | n.d. | n.d. | n.d. | n.d. | 1,85E+02 |
| 584 | WT | sham | n.d. | n.d. | n.d. | n.d. | n.d. | n.d. | n.d. |
| 585 | WT | YF 17D | 2,75E+02 | n.d. | n.d. | n.d. | n.d. | n.d. | n.d. |
| 586 | WT | YF 17D | 3,26E+02 | n.d. | n.d. | n.d. | n.d. | n.d. | n.d. |
| 587 | WT | YF 17D | 1,93E+03 | n.d. | 6,24E+02 | n.d. | n.d. | 8,36E+02 | 1,25E+02 |
| 588 | WT | YF 17D | n.d. | n.d. | n.d. | n.d. | n.d. | n.d. | n.d. |
| 589 | WT | YF 17D | 1,66E+03 | n.d. | n.d. | n.d. | n.d. | n.d. | n.d. |
| 590 | WT | YF 17D | n.d. | n.d. | n.d. | n.d. | n.d. | n.d. | n.d. |
| 591 | WT | YF-S0 | n.d. | n.d. | n.d. | n.d. | n.d. | n.d. | n.d. |
| 592 | WT | YF-S0 | n.d. | n.d. | n.d. | n.d. | n.d. | n.d. | 1,14E+02 |
| 593 | WT | YF-S0 | n.d. | n.d. | n.d. | n.d. | n.d. | n.d. | n.d. |
| 594 | WT | YF-S0 | n.d. | n.d. | n.d. | n.d. | n.d. | n.d. | n.d. |
| 595 | WT | YF-S0 | n.d. | n.d. | n.d. | 1,86E+02 | n.d. | n.d. | n.d. |
| 596 | WT | YF-S0 | n.d. | n.d. | n.d. | 1,73E+02 | n.d. | n.d. | n.d. |
| 597 | STAT2-/- | YF 17D | 1,85E+06 | 1,92E+05 | 9,94E+07 | 9,41E+06 | 3,12E+07 | 2,15E+05 | 6,31E+06 |
| 598 | STAT2-/- | YF 17D | 5,23E+07 | 2,59E+06 | 1,49E+08 | 1,91E+07 | 4,80E+07 | n.d. | 1,65E+07 |

1. Limit of detection (LoD) is around 44 copies per mg tissue, values below marked as non-detectable (n.d.)
2. Values below LoD represented as the square root of the LoD on the figure

**Supplementary Table 2. (Related to Fig. 2B, Serum)**

| Sample | Genotype | vaccine | Day 1 | Day 2 | Day 3 | Day 4 | Day 5 | Day 7 | Day 9 | Day 11 | Day 15 | Day 22 | Day 29 |
|---|---|---|---|---|---|---|---|---|---|---|---|---|---|
| 165 | WT | sham | n.d. | n.d. | n.d. | n.d. | n.d. | n.d. | n.d. | n.d. | n.d. | N/A | n.d. |
| 166 | WT | sham | n.d. | n.d. | n.d. | n.d. | n.d. | n.d. | n.d. | n.d. | n.d. | N/A | n.d. |
| 167 | WT | sham | n.d. | n.d. | n.d. | n.d. | n.d. | n.d. | n.d. | n.d. | n.d. | N/A | n.d. |
| 168 | WT | sham | n.d. | n.d. | n.d. | n.d. | n.d. | n.d. | n.d. | n.d. | n.d. | N/A | n.d. |
| 169 | WT | YF 17D | 4,87E+04 | n.d. | n.d. | n.d. | n.d. | n.d. | n.d. | n.d. | n.d. | N/A | n.d. |
| 170 | WT | YF 17D | n.d. | 1,61E+05 | n.d. | n.d. | n.d. | n.d. | n.d. | n.d. | n.d. | N/A | n.d. |
| 171 | WT | YF 17D | 4,48E+04 | 2,54E+04 | 9,55E+04 | n.d. | n.d. | n.d. | n.d. | n.d. | n.d. | N/A | n.d. |
| 172 | WT | YF 17D | n.d. | 4,81E+04 | 1,48E+05 | 1,39E+04 | N/A | n.d. | n.d. | n.d. | n.d. | N/A | n.d. |
| 173 | WT | YF 17D | 2,96E+04 | 4,34E+04 | 1,70E+05 | 1,43E+04 | n.d. | n.d. | n.d. | n.d. | n.d. | N/A | n.d. |
| 174 | WT | YF 17D | n.d. | 4,73E+04 | 9,20E+04 | N/A | n.d. | n.d. | n.d. | n.d. | n.d. | N/A | n.d. |
| 831 | WT | YF-S0 | n.d. | n.d. | 4,30E+04 | n.d. | n.d. | n.d. | n.d. | n.d. | n.d. | N/A | n.d. |
| 832 | WT | YF-S0 | n.d. | n.d. | n.d. | n.d. | n.d. | n.d. | n.d. | n.d. | n.d. | N/A | n.d. |
| 833 | WT | YF-S0 | n.d. | n.d. | n.d. | n.d. | n.d. | n.d. | n.d. | n.d. | n.d. | N/A | n.d. |
| 834 | WT | YF-S0 | n.d. | n.d. | n.d. | n.d. | n.d. | n.d. | n.d. | n.d. | n.d. | N/A | n.d. |
| 835 | WT | YF-S0 | n.d. | n.d. | n.d. | n.d. | n.d. | n.d. | n.d. | n.d. | n.d. | N/A | n.d. |
| 836 | WT | YF-S0 | n.d. | n.d. | n.d. | n.d. | n.d. | n.d. | n.d. | n.d. | n.d. | N/A | n.d. |
| 837 | STAT2-/- | YF 17D | 6,16E+04 | 1,53E+06 | 2,43E+07 | 1,40E+07 | 3,06E+06 | 6,69E+06 | | | | | |
| 838 | STAT2-/- | YF 17D | N/A | 1,61E+06 | 3,41E+07 | 1,24E+07 | 1,72E+06 | 1,24E+07 | | | | | |
| 839 | STAT2-/- | YF 17D | n.d. | 1,60E+06 | 1,38E+07 | 9,89E+06 | 2,01E+06 | 7,92E+06 | | | | | |

1. Limit of detection (LoD) is around 8900 copies per ml serum, values below marked as non-detectable (n.d.)
2. Values below LoD represented as the square root of the LoD on the figure

**Supplementary Table 3. (Related to Fig. 2C, Urine)**

| Sample | Genotype | vaccine | Day 1 | Day 2 | Day 3 | Day 4 | Day 5 | Day 7 | Day 9 | Day 11 | Day 15 | Day 22 | Day 29 |
|---|---|---|---|---|---|---|---|---|---|---|---|---|---|
| 165 | WT | sham | n.d. | n.d. | n.d. | n.d. | n.d. | n.d. | n.d. | N/A | n.d. | N/A | N/A |
| 166 | WT | sham | n.d. | n.d. | n.d. | n.d. | n.d. | n.d. | n.d. | N/A | n.d. | N/A | N/A |
| 167 | WT | sham | n.d. | n.d. | n.d. | n.d. | n.d. | n.d. | n.d. | n.d. | n.d. | N/A | n.d. |
| 168 | WT | sham | n.d. | n.d. | n.d. | n.d. | N/A | n.d. | n.d. | n.d. | n.d. | n.d. | n.d. |
| 169 | WT | YF 17D | n.d. | n.d. | N/A | n.d. | n.d. | n.d. | n.d. | n.d. | n.d. | N/A | n.d. |
| 170 | WT | YF 17D | n.d. | n.d. | N/A | n.d. | 1,13E+04 | n.d. | n.d. | n.d. | n.d. | n.d. | n.d. |
| 171 | WT | YF 17D | n.d. | n.d. | n.d. | n.d. | n.d. | n.d. | n.d. | n.d. | n.d. | n.d. | N/A |
| 172 | WT | YF 17D | n.d. | n.d. | n.d. | n.d. | n.d. | n.d. | n.d. | N/A | n.d. | n.d. | N/A |
| 173 | WT | YF 17D | n.d. | n.d. | n.d. | n.d. | n.d. | n.d. | n.d. | n.d. | n.d. | N/A | N/A |
| 174 | WT | YF 17D | n.d. | n.d. | n.d. | n.d. | n.d. | n.d. | N/A | n.d. | n.d. | N/A | n.d. |
| 831 | WT | YF-S0 | n.d. | n.d. | n.d. | n.d. | n.d. | n.d. | n.d. | n.d. | n.d. | n.d. | n.d. |
| 832 | WT | YF-S0 | n.d. | n.d. | n.d. | n.d. | n.d. | n.d. | n.d. | 2,56E+04 | n.d. | n.d. | N/A |
| 833 | WT | YF-S0 | n.d. | N/A | N/A | n.d. | n.d. | n.d. | n.d. | n.d. | n.d. | n.d. | N/A |
| 834 | WT | YF-S0 | n.d. | N/A | N/A | n.d. | n.d. | n.d. | n.d. | 1,35E+04 | n.d. | n.d. | n.d. |
| 835 | WT | YF-S0 | n.d. | n.d. | n.d. | n.d. | n.d. | 1,08E+06 | n.d. | n.d. | n.d. | n.d. | n.d. |
| 836 | WT | YF-S0 | n.d. | n.d. | n.d. | n.d. | N/A | n.d. | n.d. | n.d. | n.d. | N/A | n.d. |
| 837 | STAT2-/- | YF 17D | n.d. | n.d. | n.d. | 1,30E+08 | N/A | n.d. | | | | | |
| 838 | STAT2-/- | YF 17D | n.d. | 3,11E+05 | 1,60E+05 | 1,21E+08 | N/A | 1,81E+06 | | | | | |
| 839 | STAT2-/- | YF 17D | N/A | n.d. | n.d. | 7,62E+05 | N/A | n.d. | | | | | |

**1. Limit of detection (LoD) is around 8900 copies per ml urine, values below marked as non-detectable (n.d.)**
**2. Values below LoD represented as the square root of the LoD on the figure**

**Supplementary Table 4. (Related to Fig. 2D, Faeces)**

| Sample | Genotype | vaccine | Day 1 | Day 2 | Day 3 | Day 4 | Day 5 | Day 7 | Day 9 | Day 11 | Day 15 | Day 22 | Day 29 |
|---|---|---|---|---|---|---|---|---|---|---|---|---|---|
| 165 | WT | sham | n.d. | n.d. | n.d. | n.d. | n.d. | n.d. | n.d. | n.d. | n.d. | n.d. | n.d. |
| 166 | WT | sham | n.d. | n.d. | n.d. | n.d. | n.d. | n.d. | n.d. | n.d. | n.d. | n.d. | n.d. |
| 167 | WT | sham | n.d. | n.d. | n.d. | n.d. | n.d. | n.d. | n.d. | n.d. | n.d. | n.d. | n.d. |
| 168 | WT | sham | n.d. | n.d. | n.d. | n.d. | n.d. | n.d. | n.d. | n.d. | n.d. | n.d. | n.d. |
| 169 | WT | YF 17D | n.d. | n.d. | n.d. | 1,04E+05 | n.d. | n.d. | n.d. | n.d. | n.d. | n.d. | n.d. |
| 170 | WT | YF 17D | n.d. | 5,00E+04 | n.d. | n.d. | n.d. | n.d. | n.d. | n.d. | n.d. | n.d. | n.d. |
| 171 | WT | YF 17D | n.d. | n.d. | n.d. | n.d. | n.d. | n.d. | n.d. | n.d. | n.d. | n.d. | n.d. |
| 172 | WT | YF 17D | n.d. | N/A | n.d. | n.d. | n.d. | n.d. | n.d. | n.d. | n.d. | n.d. | n.d. |
| 173 | WT | YF 17D | n.d. | n.d. | n.d. | n.d. | n.d. | n.d. | n.d. | n.d. | n.d. | n.d. | n.d. |
| 174 | WT | YF 17D | n.d. | n.d. | n.d. | n.d. | n.d. | n.d. | n.d. | n.d. | n.d. | n.d. | n.d. |
| 831 | WT | YF-S0 | n.d. | n.d. | n.d. | n.d. | n.d. | n.d. | n.d. | n.d. | n.d. | n.d. | n.d. |
| 832 | WT | YF-S0 | n.d. | n.d. | n.d. | n.d. | n.d. | n.d. | n.d. | n.d. | n.d. | n.d. | n.d. |
| 833 | WT | YF-S0 | n.d. | n.d. | n.d. | n.d. | n.d. | n.d. | n.d. | n.d. | n.d. | n.d. | n.d. |
| 834 | WT | YF-S0 | n.d. | n.d. | n.d. | n.d. | n.d. | n.d. | n.d. | n.d. | n.d. | n.d. | n.d. |
| 835 | WT | YF-S0 | n.d. | n.d. | n.d. | n.d. | n.d. | n.d. | n.d. | n.d. | n.d. | n.d. | n.d. |
| 836 | WT | YF-S0 | n.d. | n.d. | n.d. | 6,35E+04 | n.d. | n.d. | n.d. | n.d. | n.d. | n.d. | n.d. |
| 837 | STAT2-/- | YF 17D | n.d. | n.d. | 2,17E+04 | n.d. | 5,93E+07 | 1,25E+07 | | | | | |
| 838 | STAT2-/- | YF 17D | n.d. | n.d. | n.d. | 5,96E+06 | n.d. | 1,52E+05 | | | | | |
| 839 | STAT2-/- | YF 17D | n.d. | n.d. | n.d. | 7,28E+06 | n.d. | 2,40E+05 | | | | | |

1. Limit of detection (LoD) is around 1500 copies per ml faeces, values below marked as non-detectable (n.d.)
2. Values below LoD represented as the square root of the LoD on the figure

**Supplementary Table 5. (Related to Fig. 2E, Buccal swab)**

| Sample | Genotype | vaccine | Day 1 | Day 2 | Day 3 | Day 4 | Day 5 | Day 7 | Day 9 | Day 11 | Day 15 | Day 22 | Day 29 |
|---|---|---|---|---|---|---|---|---|---|---|---|---|---|
| 165 | WT | sham | n.d. | n.d. | n.d. | n.d. | n.d. | n.d. | n.d. | n.d. | n.d. | n.d. | n.d. |
| 166 | WT | sham | n.d. | n.d. | n.d. | n.d. | n.d. | n.d. | n.d. | n.d. | n.d. | n.d. | n.d. |
| 167 | WT | sham | n.d. | n.d. | n.d. | n.d. | n.d. | n.d. | n.d. | n.d. | n.d. | n.d. | n.d. |
| 168 | WT | sham | n.d. | n.d. | n.d. | n.d. | n.d. | n.d. | n.d. | n.d. | n.d. | n.d. | n.d. |
| 169 | WT | YF 17D | n.d. | n.d. | n.d. | n.d. | n.d. | n.d. | n.d. | n.d. | n.d. | n.d. | n.d. |
| 170 | WT | YF 17D | n.d. | n.d. | n.d. | n.d. | n.d. | n.d. | n.d. | n.d. | n.d. | n.d. | n.d. |
| 171 | WT | YF 17D | n.d. | n.d. | n.d. | n.d. | n.d. | n.d. | n.d. | n.d. | n.d. | n.d. | n.d. |
| 172 | WT | YF 17D | n.d. | n.d. | n.d. | n.d. | n.d. | n.d. | n.d. | n.d. | n.d. | n.d. | n.d. |
| 173 | WT | YF 17D | n.d. | n.d. | n.d. | n.d. | n.d. | n.d. | n.d. | n.d. | n.d. | n.d. | n.d. |
| 174 | WT | YF 17D | 3,98E+03 | n.d. | n.d. | n.d. | n.d. | n.d. | n.d. | n.d. | n.d. | n.d. | n.d. |
| 831 | WT | YF-S0 | n.d. | n.d. | n.d. | 5,08E+03 | n.d. | n.d. | n.d. | n.d. | n.d. | n.d. | n.d. |
| 832 | WT | YF-S0 | n.d. | n.d. | n.d. | n.d. | 1,66E+03 | n.d. | n.d. | n.d. | n.d. | n.d. | n.d. |
| 833 | WT | YF-S0 | n.d. | n.d. | n.d. | n.d. | n.d. | n.d. | n.d. | n.d. | n.d. | n.d. | n.d. |
| 834 | WT | YF-S0 | n.d. | n.d. | n.d. | n.d. | n.d. | n.d. | n.d. | n.d. | n.d. | n.d. | n.d. |
| 835 | WT | YF-S0 | n.d. | n.d. | n.d. | n.d. | n.d. | n.d. | n.d. | n.d. | n.d. | n.d. | n.d. |
| 836 | WT | YF-S0 | n.d. | n.d. | n.d. | n.d. | 3,81E+03 | n.d. | n.d. | n.d. | n.d. | n.d. | n.d. |
| 837 | STAT2-/- | YF 17D | n.d. | n.d. | 1,28E+05 | 8,60E+05 | 2,88E+06 | 3,33E+07 | | | | | |
| 838 | STAT2-/- | YF 17D | n.d. | n.d. | 2,07E+05 | 7,92E+05 | n.d. | 2,21E+06 | | | | | |
| 839 | STAT2-/- | YF 17D | n.d. | n.d. | 4,29E+04 | 8,95E+05 | 3,16E+07 | 1,34E+07 | | | | | |

**1. Limit of detection (LoD) is around 1500 copies per ml buccal swab, values below marked as non-detectable (n.d.)**
**2. Values below LoD represented as the square root of the LoD on the figure**
